# PipeIT2: A tumor-only somatic variant calling workflow for Molecular Diagnostic Ion Torrent sequencing data

**DOI:** 10.1101/2022.06.28.497937

**Authors:** Desiree Schnidrig, Andrea Garofoli, Andrej Benjak, Gunnar Rätsch, Mark A. Rubin, SOCIBP consortium, Salvatore Piscuoglio, Charlotte K. Y. Ng

**Affiliations:** Department for BioMedical Research, University of Bern, Bern, Switzerland; SIB Swiss Institute of Bioinformatics, Lausanne, Switzerland; Institute of Medical Genetics and Pathology, University Hospital Basel, University of Basel, Basel, Switzerland; Department of Computer Science, ETH Zurich; Bern Center for Precision Medicine, Bern, Switzerland; Department of Biomedicine, University Hospital Basel, University of Basel, Basel, Switzerland

**Author notes:** **Correspondence:** Dr. Charlotte K. Y. Ng. Department for BioMedical Research, University of Bern, Murtenstrasse 40, Bern, 3008, Switzerland. Co-first authors (Desiree Schnidrig, Andrea Garofoli). **Disclosures:** C.K.Y.N. is a consultant for Repare Therapeutics.

## Abstract

Precision oncology relies on the accurate identification of somatic mutations in cancer patients. While the sequencing of the tumoral tissue is frequently part of routine clinical care, the healthy counterparts are rarely sequenced. We previously published PipeIT, a somatic variant calling workflow specific for Ion Torrent sequencing data enclosed in a Singularity container. PipeIT combines user-friendly execution, reproducibility and reliable mutation identification, but relies on matched germline sequencing data to exclude germline variants. Expanding on the original PipeIT, here we describe PipeIT2 to address the clinical need to define somatic mutations in the absence of germline control. We show that PipeIT2 achieves a >95% recall for variants with variant allele fraction >10%, reliably detects driver and actionable mutations and filters out most of the germline mutations and sequencing artifacts. With its performance, reproducibility and ease of execution, PipeIT2 is a valuable addition to molecular diagnostics laboratories.

## INTRODUCTION

Detection of genomic alterations is becoming a critical component in the standard-of-care in modern oncology^1,2^. Typically, the detection of genomic alterations is performed using targeted sequencing panels to profile previously described cancer and actionable gene regions. The Ion Torrent sequencing platform is frequently used for targeted sequencing in the diagnostic setting due to its relatively low costs, ability to profile limited genetic material and rapid turnaround^3^. While Ion Torrent library preparation and sequencing are relatively straightforward, the methods for sequencing data analysis are not very well-developed. Due to the technical differences between Ion Torrent and other sequencing platforms, most of the variant calling tools previously tested, validated and extensively used by the community are not suited for Ion Torrent data. Ion Torrent sequencing data are typically analyzed on its own analysis platform Ion Reporter. We and others have reported the high false positive rate of Ion Reporter analyses, especially for custom panels that lack built-in analysis workflows^4,5^. Consequently, analyses performed on the Ion Reporter platform typically require extensive manual review of the results.

We recently published PipeIT, a pipeline to detect somatic variants in matched tumor-germline samples from Ion Torrent sequencing data^5^, providing a reliable and automated workflow to perform variant calling analysis, outperforming a standard Ion Reporter analysis. We previously benchmarked the variant calling analysis of Ion Reporter using both standard parameters provided by the manufacturer and a set of optimized parameters. In both cases, Ion Reporter was indeed able to detect genuine somatic mutations, validated by whole-exome sequencing and/or Sanger sequencing on two different matched tumor-germline cohorts), but it also showed the presence of several false positives, notably when the analysis was performed using the standard, non optimized parameters provided by the machine^5^. To ensure reproducibility and ease of deployment, PipeIT was built as a Singularity^6^ container image file that can be easily executed with a single command, without the need of additional software other than the Singularity platform.

The main drawback of PipeIT is the need for germline matched control data. When the goal is to identify somatic mutations, the sequencing of normal controls can be critical in order to remove germline mutations^1,7,8^. In routine clinical care, however, the sequencing of tumor-only tissue is often preferred, for time, costs and sample availability reasons. Moreover, researchers might want to analyze old, archived samples, for which matched germline controls may not be available. These scenarios significantly limit the contexts where PipeIT can be used and, ultimately, prevent the software from fully achieving its original aim.

Here we present PipeIT2, an extension of PipeIT to enable variant calling analyses on tumor samples without matched germline controls with a single command. PipeIT2 identifies and filters likely germline mutations by leveraging their allele frequencies in population databases and, if provided, by detecting their presence in unmatched Panel of Normal (PoN) samples. We demonstrate that PipeIT2 was able to detect clinically relevant somatic mutations, while correctly identifying and removing most of the germline genomic alterations.

## MATERIALS AND METHODS

### Building the PipeIT2 Singularity Container Image

The original PipeIT Singularity container has been updated to include the PipeIT2 tumor-only workflow. The file is a read-only squashfs file system Singularity image built on a CentOS7 Docker image as a base, as previously described^5^. PipeIT2 provides the entry points to perform both the matched tumor-germline and the new tumor-only workflow. Similar to PipeIT, the new PipeIT2 Singularity image provides most of the data needed to perform the complete analysis, except the population datasets due to file size. The population datasets can be downloaded with PipeIT2 using a utility provided in the Singularity image.

### The PipeIT2 tumor-only analysis workflow

The PipeIT2 tumor-only analysis workflow comprises the following steps: 1) variant calling, 2) variant post-processing, 3) variant annotation, 4) read count and quality-based variant filtering, 5) annotation-based variant filtering and, 6) optionally, PoN-based variant filtering (**Figure 1**). Due to their likely role in cancer development, hotspot variants are annotated and whitelisted from all filtering steps^9,10^. This workflow requires a Binary Alignment Map (BAM)^11^ file for the tumor sample from the Ion Torrent Server aligned using the Torrent Mapping Alignment Program (TMAP) aligner, a Browser Extensible Data (BED)^12^ file defining the target sequenced regions, Annovar^13^ annotation files comprising of population minor allele frequencies, and optionally blacklist BED file and/or a Variant Call Format (VCF)^14^ file containing the mutations found in the PoN. In contrast to the original PipeIT tumor-germline analysis workflow, PipeIT2 does not use sequencing data from matched germline controls.

**Figure 1.**
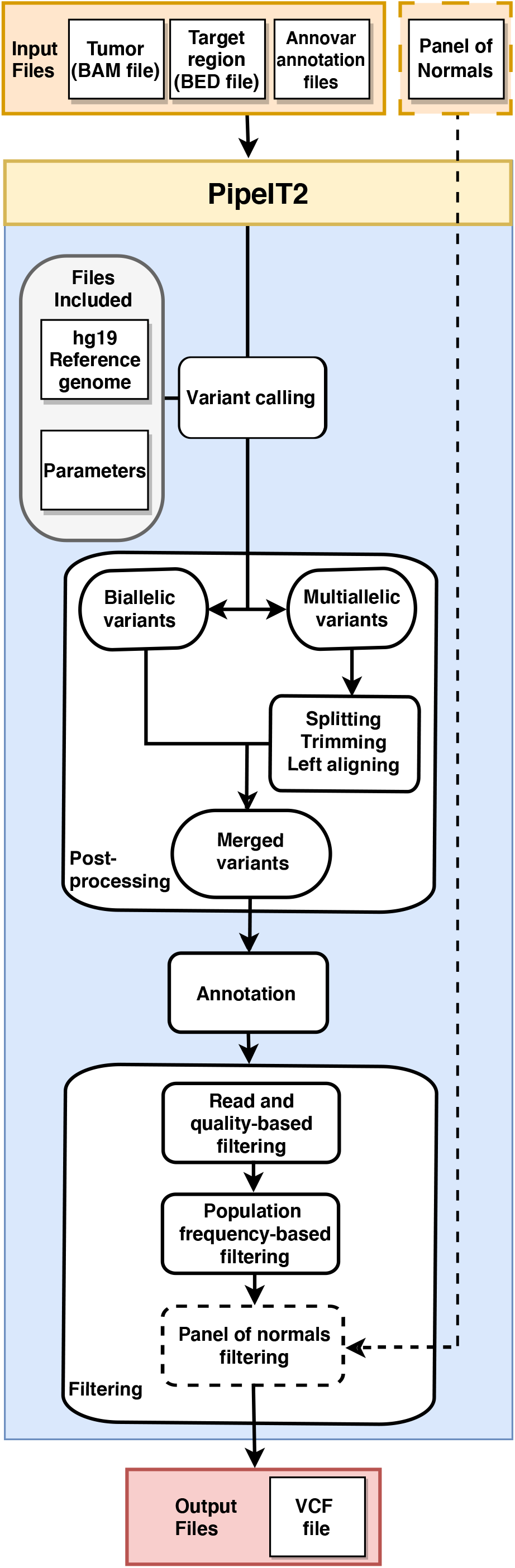
Overview of the PipeIT2 tumor-only workflow. Flowchart showing the steps of the workflow. The workflow takes the BAM file for the tumor sample, the BED file for the target regions, the Annovar datasets for the population databases and, optionally, a Panel of Normals. Variant calling is then performed using the Torrent Variant Caller with the packaged parameters file. Mutations are filtered based on read count and quality, population frequencies and, when provided, the Panel of Normals. The output is returned as a VCF file.

Variant calling (step 1) is performed using the Torrent Variant Caller (TVC, v5.12-27 with tvcutils 5.0-3, Thermo Fisher Scientific) using the same low stringency parameters used in the original PipeIT tumor-germline analysis workflow^5^, packaged in a JSON file within PipeIT2. Specifically, we use a quality threshold of 6.5, a variant score equal or higher than 10, a minimum coverage of 8 reads for single nucleotide variants (SNVs) and 15 reads for small insertion/deletions (INDELs) and a variant allele frequency (VAF) of 2% for both SNVs and INDELs. It is possible to customize the parameters by providing PipeIT2 a JSON file following the format required by TVC. Some commercially available gene panels come with a blacklist, consisting of recurrent artifacts identified through the sequencing of normal samples. The blacklist is typically included in the hotspot BED file and these variants are tagged with “BSTRAND=F” (on the forward strand), “BSTRAND=R” (on the reverse strand), or “BSTRAND=B” (on both strands). If a blacklist BED file is provided, it will be used by TVC. Normalization of the raw variants (step 2, splitting multiallelic into biallelic variants and left-aligning) is then performed as in PipeIT to facilitate downstream processing.

In the next step, normalized variants are annotated using snpEff^15^ and Annovar^13^ (step 3). Aside from the transcript and protein effects of the variants, PipeIT2 also annotates the variants with their homopolymer lengths and their minor allele frequencies observed in populations using data from the 1000 Genomes Project (1KG)^16^, the Exome Aggregation Consortium (ExAC)^17^, the NHLBI Exome Sequencing Project (ESP)^18^ and the Genome Aggregation Database (GnomAD)^19^. Additionally, variants in mutation hotspot regions^9,10^ [https://github.com/charlottekyng/cancer_hotspots, last accessed March 9, 2022] are annotated.

Variant filtering is then performed in three stages. First, read count and quality-based filtering (step 4) is performed to remove variants of low confidence. By default, PipeIT2 removes variants with fewer than 20 total reads (corresponding to the INFO field FDP), fewer than 8 reads supporting the variant (FAO), less than 10% VAF (FAO/FDP), fewer than 3 forward (FSAF) and 3 reverse reads (FSAR), strand bias (FSAF/FSAR) below 0.2 in either direction, a quality score below 15, or variants in homopolymer regions of length greater than 4 (**Table 1**).

**Table 1.** Filtering parameters and default values of the tumor-only workflow.

| Parameter | Description | Default value |
| --- | --- | --- |
| --min_supporting_reads | Minimum number of reads supporting the variant | 8 |
| --min_tumor_depth | Minimum read depth at the locus | 20 |
| --min_allele_fraction | Minimum allele fraction (i.e. the number of read supporting the variant divided by the read depth at the locus) | 0.1 |
| --homopolymer_run | Maximum homopolymer region length | 4 |
| --max_pop_af | Maximum frequency of mutation in population databases | 0.005 |
| --quality | Minimum quality score | 15 |

Second, PipeIT2 leverages population data to remove likely germline variants (step 5). Specifically, variants are removed if they are observed with minor allele frequencies equal to or higher than 0.5% in any of the four population-level databases 1KG, ExAC, ESP and GnomAD. Variants with VAF between 0.4 and 0.6, or greater than 0.9 are removed if they are found at any allele frequency in any of the four population-level datasets.

Third, as an optional step, PipeIT2 can use a user-defined Panel of Normals (PoN) in order to further reduce the number of likely false positive variants (step 6), including germline variants not removed in step 5 and systematic sequencing and alignment artifacts. Accepted inputs are either a pre-generated PoN VCF file or a list of unmatched germline BAM files from samples sequenced on the same platform as the tumor sample. If a list of BAM files is provided, PipeIT2 automatically calls variants in each of these normal samples as per variant calling and post-processing steps in the tumor-only workflow. These germline VCF files are then merged with the GATK ‘CombineVariants’ function using the UNIQUIFY option and retaining mutations found in at least two of the input samples.

The final post-filtering output is returned as a VCF file.

### Evaluation of the PipeIT2 tumor-only workflow

Sequencing data from 15 formalin-fixed paraffin-embedded colon adenomas^20^ (COAD cohort) and 10 frozen hepatocellular carcinoma samples^21^ (HCC cohort) were retrieved from our previous publication^5^. The performance of the PipeIT2 tumor-only workflow and the contribution of the PoN-based variant filtering step (step 6 above) was assessed using the outputs from the tumor-germline workflow as the benchmark. The PoN files used in these analyses were generated from 8 randomly selected unmatched germline samples from the corresponding cohorts. The mutations detected in PipeIT2 were classified as: true positives (TP, mutations called by both workflows), false positives (FP, mutations called by the tumor-only workflow but not the tumor-germline workflow), and false negatives (FN, mutations detected by the tumor-germline workflow but not the tumor-only workflow). Performance of PipeIT2 was evaluated as recall (TP/(TP+FN)), precision (TP/(TP+FP)) and F1 score (2*precision*recall/(precision+recall)).

### Visualization of BAM files

Integrative Genomics Viewer (IGV)^22^ was used to visualize the BAM files and search for the presence of false positive mutations across the original matched tumor-germline pairs and the unmatched germline samples used to build the PoN files for these benchmarking analyses.

## SOFTWARE AVAILABILITY

PipeIT2 is available at https://github.com/ckynlab/PipeIT2.

## RESULTS

### Running the PipeIT2 tumor-only workflow

To provide an effective somatic variant calling analysis on tumor data originated from Ion Torrent platform in the absence of a matched germline, we updated the original PipeIT functionality to allow the users to choose between the classic tumor-germline (PipeIT) and the new tumor-only (PipeIT2) analyses. The PipeIT2 tumor-only workflow (**Figure 1**) can be executed in a single command as follows:

singularity run PipeIT.img -t path/to/tumor.bam -e path/to/region.bed -c path/to/annovar/humandb/folder (-d path/to/PoN/file.vcf)

Using this command, somatic variants are called with an Ion Torrent-specific variant caller (TVC), followed by a normalization step to facilitate downstream processing. Raw variant calls are filtered in a multi-step process, specifically optimized to remove likely germline and artefactual variants in the absence of a matched germline control. Specifically, low confidence variants are removed with read- and quality-based filters. Then, information from population sequencing data is leveraged to identify likely germline variants. An optional panel of unmatched normal samples (PoN) can be used to further reduce the number of germline and artefactual variants. In order to ensure the detection of known cancer hotspot variants, they are annotated and whitelisted from all filtering steps^9,10^.

### Evaluation of the PipeIT2 tumor-only workflow

To evaluate the performance of the PipeIT2 tumor-only workflow, we analyzed the 10 fresh frozen hepatocellular carcinoma (HCC) samples and 15 formalin-fixed paraffin-embedded colon adenomas (COAD) used in our previous publication^5^. The 10 HCCs and their matched germline were sequenced using a previously published custom HCC targeted sequencing panel^21^ and the 15 COADs with corresponding germline samples using the Oncomine Comprehensive Panel v3^23^. We ran the tumor-only workflow with default parameters (**Table 1**) to call somatic variants and compared the non-synonymous and *TERT* promoter mutations to those called using the tumor-germline workflow. To investigate whether the use of a PoN could improve the performance, for each of the 25 samples, a PoN VCF was generated from 8 randomly chosen unmatched germline samples (i.e. excluding the matched germline) of the corresponding cohort. We analyzed each of these 25 samples with and without the PoN and evaluated the performance of the tumor-only workflow in terms of precision, recall and F1 value.

Across the 10 HCC samples, we identified 53 true positive, 11 false positive and 15 false negative variants (**Figure 2A**). Of the 53 true positive variants, 10 were annotated hotspot variants. All 11 false positive variants were confirmed as rare germline variants on IGV (**Supplemental Figures 1 and 2**). Nine of them are the same recurring dinucleotide variant (DNV) *chr2:21232803:TG>CA* in *APOB*, which upon closer inspection appeared to be 2 distinct SNPs - rs584542 (*chr2:21232803:T>C*) and rs1041968 (*chr2:21232804:G>A*) which were validated as germline by orthogonal whole-exome sequencing^21^ (**Supplemental Figure 2**). This variant was also present in the PoN and therefore successfully filtered out in the PoN analysis (**Supplemental Figure 2**). All 15 false negative variants were removed by filters specific to the tumor-only workflow to limit the number of artifactual variants. In particular, 14 variants were below the VAF filtering threshold of 10% and one variant was located in a homopolymer region of length greater than 4. It is worth mentioning that one of the HCC samples (HPU207) was previously identified as hypermutated^21^ and 13/15 of the false negative variants were missed in this sample. Overall, the analysis without a PoN achieved recall, precision and F1 of 0.78, 0.83 and 0.80 respectively (**Figure 2B**). With the use of a PoN, precision improved to 0.96, resulting in an F1 score of 0.86. When we only considered variants >10% VAF, a threshold typically used in the molecular diagnostic setting, the recall increased from 0.78 to 0.98 with an F1 score of 0.90 in the analysis without a PoN and 0.97 with the additional use of a PoN (**Figure 2B**).

**Figure 2.**
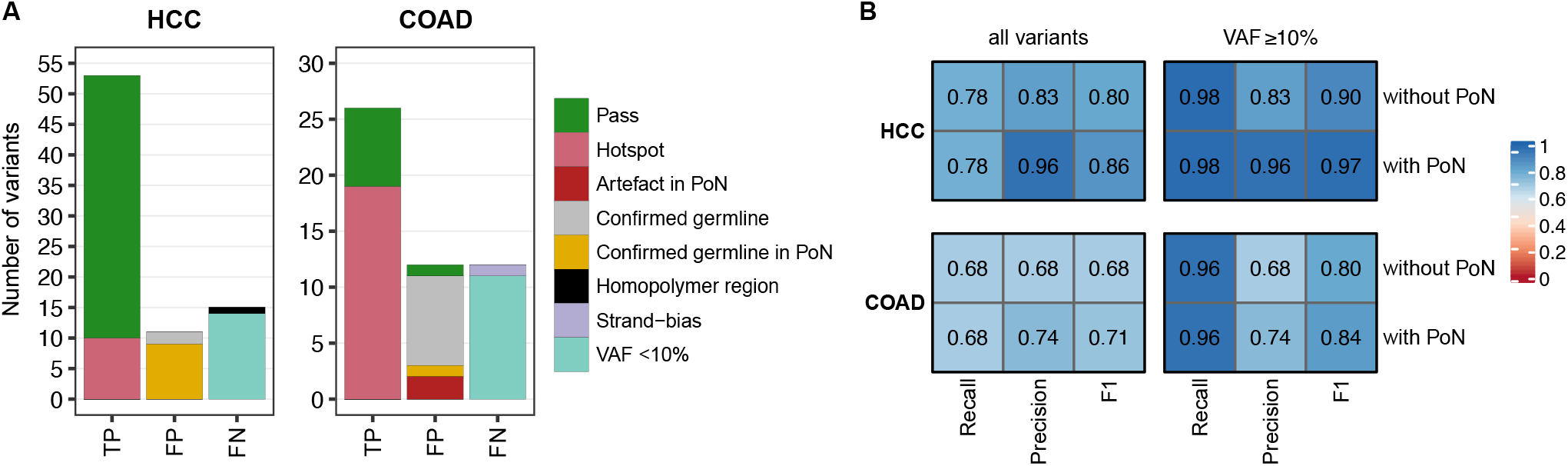
Performance evaluation of PipeIT2. **(A)** Barplots showing the number of true positive (TP), false positive (FP) and false negative (FN) variants in the (left) HCC and (right) COAD cohorts. Mutation classification is indicated in the color key. (**B)** Heatmaps showing the recall, precision and F1 of PipeIT2 in a VAF range of (left) 1%-100% (‘all variants’) and (right) 10%-100% in the (top) HCC and (bottom) COAD cohorts. Boxes are colored according to the color key.

In the cohort of 15 COADs, we identified 26 true positives, including 19 hotspot variants, as well as 12 false positive and 12 false negative variants (**Figure 2A**). Most (9/12) false positive variants were confirmed as rare germline variants, including one that was successfully removed in the PoN analysis. Another two artifactual variants were present in the respective PoNs and hence successfully filtered out in the PoN analysis. Similar to the analysis of the HCC cohort, nearly all (11/12) false negative variants were filtered out due to their low allele frequency (VAF < 10%). The remaining false negative variant was removed due to its strand-bias. Without the use of a PoN, the recall, precision and F1 score were all 0.68, while the precision increased to 0.74 (F1=0.71) with the use of a PoN (**Figure 2B**). Excluding variants with VAF< 10%, the recall was 0.96, increasing the F1 score to 0.80 and 0.84 in the analysis with and without PoN, respectively (**Figure 2B**).

Overall, PipeIT’s recall of variants with a VAF ≥10% was nearly perfect, with only one variant missed in each cohort. Misclassification of rare germline variants as somatic was the main reason for false positive variants (20/23; 87%) and represents a known limitation of tumor-only variant calling. The additional use of a PoN has helped to reduce the overall number of false positives by 52% (12/23).

### Evaluation of the PipeIT2 tumor-only workflow in a clinical context

To evaluate whether the PipeIT2 tumor-only workflow would detect clinically and biologically significant variants, we used oncoKB^24^ to annotate the oncogenicity and clinical actionability (levels 1-3, namely FDA-approved drugs, standard care and clinical evidence) of the variants. Across both cohorts, the PipeIT2 tumor-only workflow successfully detected all cancer hotspot variants. In the HCC cohort, we detected the known oncogenic *TERT* promoter (c.-150C>T) and *CTNNB1* (p.S33C; p.T41A) mutations, likely oncogenic variants in *CTNNB1* (p.D32A; p.S37C) and likely oncogenic truncating variants in *ARID1A* (p.Y128*; p.S255fs), *ATM* (p.C117*), *AXIN1* (p.Q559*), *RB1* (p.E545*) and *TP53* (p.C135*; **Figure 3A**). In the COAD cohort, PipeIT2 identified several targetable oncogenic variants such as a *KRAS* p.G12C and *BRAF* p.V600E, as well as mutations linked to anti-EGFR resistance such as the *KRAS* and *NRAS* p.Q61K variants (**Figure 3B)**. In addition, oncogenic variants in *BRAF* (p.N581I), *CTNNB1* (p.T41A; p.S45A) and *PIK3CA* (p.C420R), a likely oncogenic truncating variant in *ARID1A* (p.Y815fs) and a likely oncogenic variant in *KDR* (p.C482R) were also identified.

**Figure 3.**
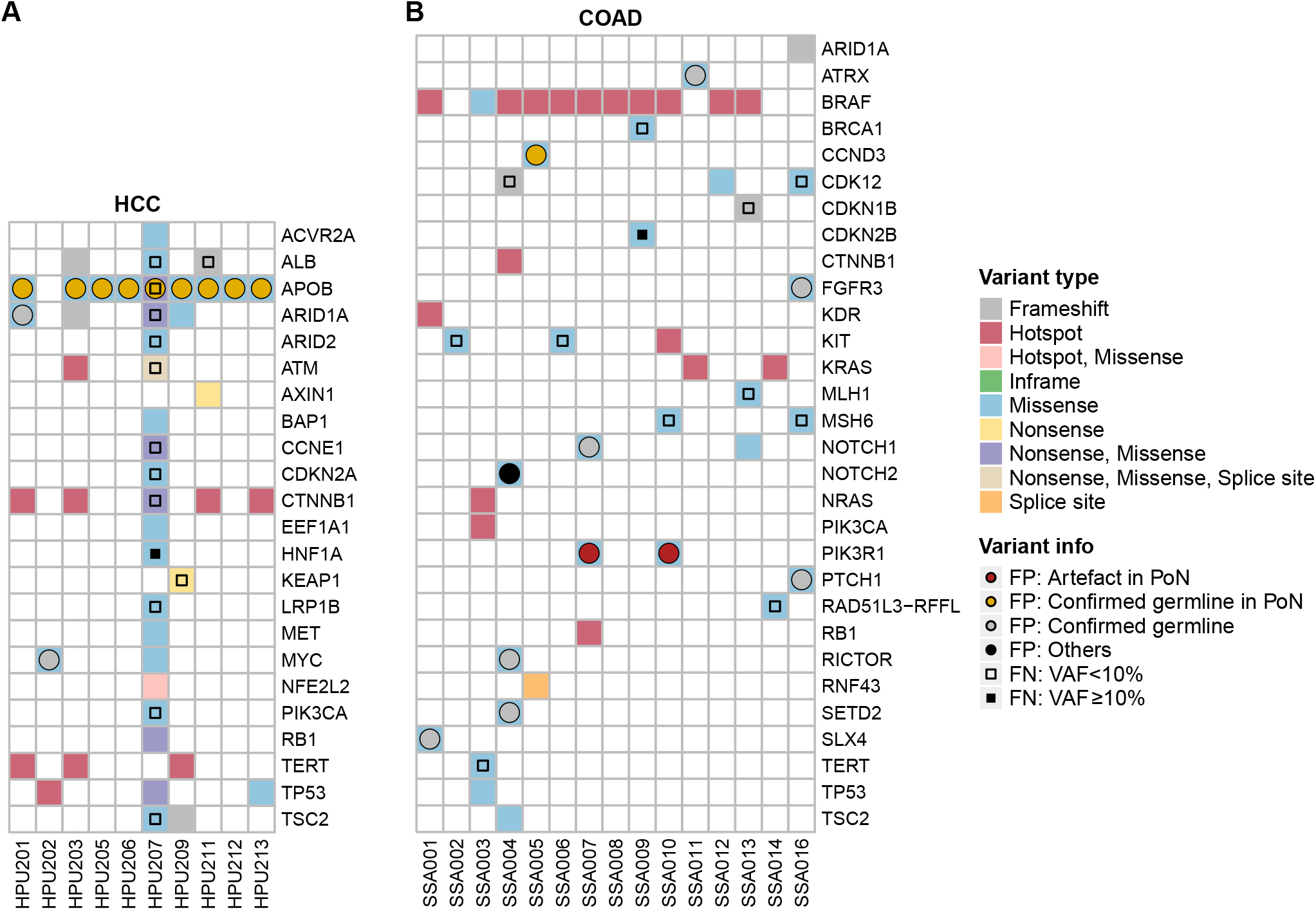
Variants detected by PipeIT2. Oncoprints of the variants called in the **(A)** HCC and **(B)** COAD cohorts. Variant types are color-coded as indicated in the color key. Multiple variant types indicate multiple variants of different types. False positive mutations are marked with a dot. Red dots indicate likely sequencing artifacts found in the PoN, yellow dots indicate confirmed germline variants found in the PoN, gray dots indicate confirmed germline variants absent in the PoN and black dots indicate other false positive mutations. False negative mutations are highlighted with an empty square if their VAF is < 10% and with a filled square if ≥10%.

Among the 23 false positive variants (11 in the HCC cohort and 12 in the COAD cohort), 20 were germline variants in genes such as *APOB* and *NOTCH2*, of which 10 were removed with the PoN (**Figure 3)**. Of the remaining three false positives, two were likely sequencing artifacts which were filtered out with the PoN and one was likely an artifact. All 27 false negative variants were low VAF variants. Of those, 25 had a VAF < 10% and the remaining two had a VAF between 10% and 15%. Only five of these low-VAF variants, *ATM* (p.E281*), *HNF1A* (p.G375fs) and *KEAP1* (p.R554*) in the HCC cohort and *CDK12* (p.S133fs) and *CDKN1B* (p.R152fs) in the COAD cohort are likely oncogenic but none of them was reported as potential resistance variant.

## DISCUSSION

Precision oncology care is increasingly reliant on the identification of somatic DNA alterations in cancer patients. DNA sequencing of tumor tissues with targeted genomic assays represents, to date, the best means to retrieve this information^25,26^. Furthermore, the additional sequencing of a healthy tissue sample from the same cancer patient is the definitive way to determine which of the genetic alterations found in the tumor tissue are likely somatic^8^.

Ion Torrent is one of the most popular sequencing platforms in the routine diagnostic setting due to its low costs and low sample input requirements, but the proprietary Ion Reporter software requires a paid license and lacks a streamlined data analysis, particularly for custom target panels. We previously developed PipeIT, a somatic variant calling workflow specific for Ion Torrent sequencing data enclosed in a Singularity image file^5^. The strength of PipeIT lies in its ease of deployment and use, reproducible results, and demonstrated accuracy. On the other hand, the need for tumor-germline matched sequencing data limits the use of PipeIT in the clinical setting where germline samples are frequently not sequenced. The main reasons for the lack of sequencing data of a matched normal sample are time, costs and sample availability. To address this shortcoming, we developed PipeIT2, a Singularity container which contains the original PipeIT tumor-germline workflow and an additional tumor-only workflow.

To overcome the challenges associated with the lack of a matched gemline control, PipeIT2 leverages three filtering steps. The first filter relies on more stringent filtering thresholds compared to those used in the tumor-germline workflow, including a VAF threshold of 10%, compared to the previous 5%, and additional strand-bias and homopolymer filters. The second makes use of data obtained from the 1KG^16^, ExAC^17^, ESP^18^ and the GnomAD^19^. Mutations detected in at least 0.5% (or any other user-defined percentage) of the samples in any of these databases are removed from the final output. The last filter is the optional PoN filter, which consists of user-defined mutations obtained from unmatched normal samples or otherwise blacklisted variants. This third step is not mandatory, to enable the use of the tumor-only workflow even if there are no unmatched germline samples available.

To evaluate the performance of PipeIT2, the mutations identified by PipeIT2 from 10 HCCs and 15 COADs were compared to the ones identified by the tumor-germline workflow. Using panels of 8 randomly chosen unmatched normal samples for each tumor sample, a total of 79 non-synonymous or *TERT* promoter mutations, including several important clinical biomarkers, were correctly detected across the two cohorts. These include targetable mutations such as *KRAS* p.G12C and *BRAF* p.V600E, several mutations implicated in anti-EGFR resistance such as the *KRAS* and *NRAS* p.Q61K variants and various known oncogenic variants in genes such as *BRAF, CTNNB1, PIK3CA* and *TERT*. Nevertheless, 27 mutations were mistakenly removed from the PipeIT2 output. The primary reason for the removal (25/27; 93%) was the low allele fraction of these mutations. This is a result of the more stringent VAF-based filtering in the tumor-only workflow which is necessary to limit the number of false positive calls in the absence of a matched germline sample. Given that clinically important resistance mechanisms typically involve recurrent hotspots and PipeIT2 actively whitelists such hotspot mutations, these mutations would still be identified even if they are found at low VAF.

By providing a variant calling analysis able to detect somatic mutations in tumor samples lacking a matched germline control, PipeIT2 offers an important improvement over the original PipeIT workflow. Thanks to filters based on population allele frequencies and variants found in panels of unmatched germline samples, PipeIT2 was able to detect most of the somatic mutations previously identified in the matched tumor-germline analysis, including several important clinical biomarkers. In conclusion, PipeIT2 offers a powerful, user friendly and easily reproducible tool specific for Ion Torrent targeted sequencing analyses.

## ACKNOWLEDGMENTS

Development of PipeIT2 was performed at the Leonhard Med platform at ETH Zurich and the sciCORE scientific computing center at University of Basel.

## AUTHOR CONTRIBUTIONS

S.P. and C.K.Y.N. conceived and supervised the study. D.S., A.G., A.B., the SOCIBP consortium and C.K.Y.N. developed the methodology. G.R and M.A.R. provided critical review of the results. D.S., A.G. and C.K.Y.N. interpreted the results and wrote the manuscript. All authors agreed to the final version of the manuscript.

Members of the SOCIBP consortium: Andrej Benjak, Andre Kahles, Charlotte K. Y. Ng, Salvatore Piscuoglio, Gunnar Rätsch, Mark A. Rubin, Desiree Schnidrig, Senija Selimovic-Hamza

## Notes

**Grants and other funding sources:** C.K.Y.N. and S.P. were supported by the Swiss Cancer Research foundation (KFS-4543-08-2018, KFS-4988-02-2020-R, respectively). The SOCIBP (Swiss Molecular Pathology Breakthrough Platform) is a driver project funded by the Swiss Personalized Health Network (SPHN). S.P. was supported by the Surgery Department of the University Hospital Basel and by The Prof. Dr. Max Cloëtta foundation. The funders had no role in study design, data collection, and analysis, decision to publish, or preparation of the manuscript.

