## Supplementary figures and images for "PipeIT2: A tumor-only somatic variant calling workflow for Molecular Diagnostic Ion Torrent sequencing data"

### Supplemental Figure 1

Supplemental Figure 1

A

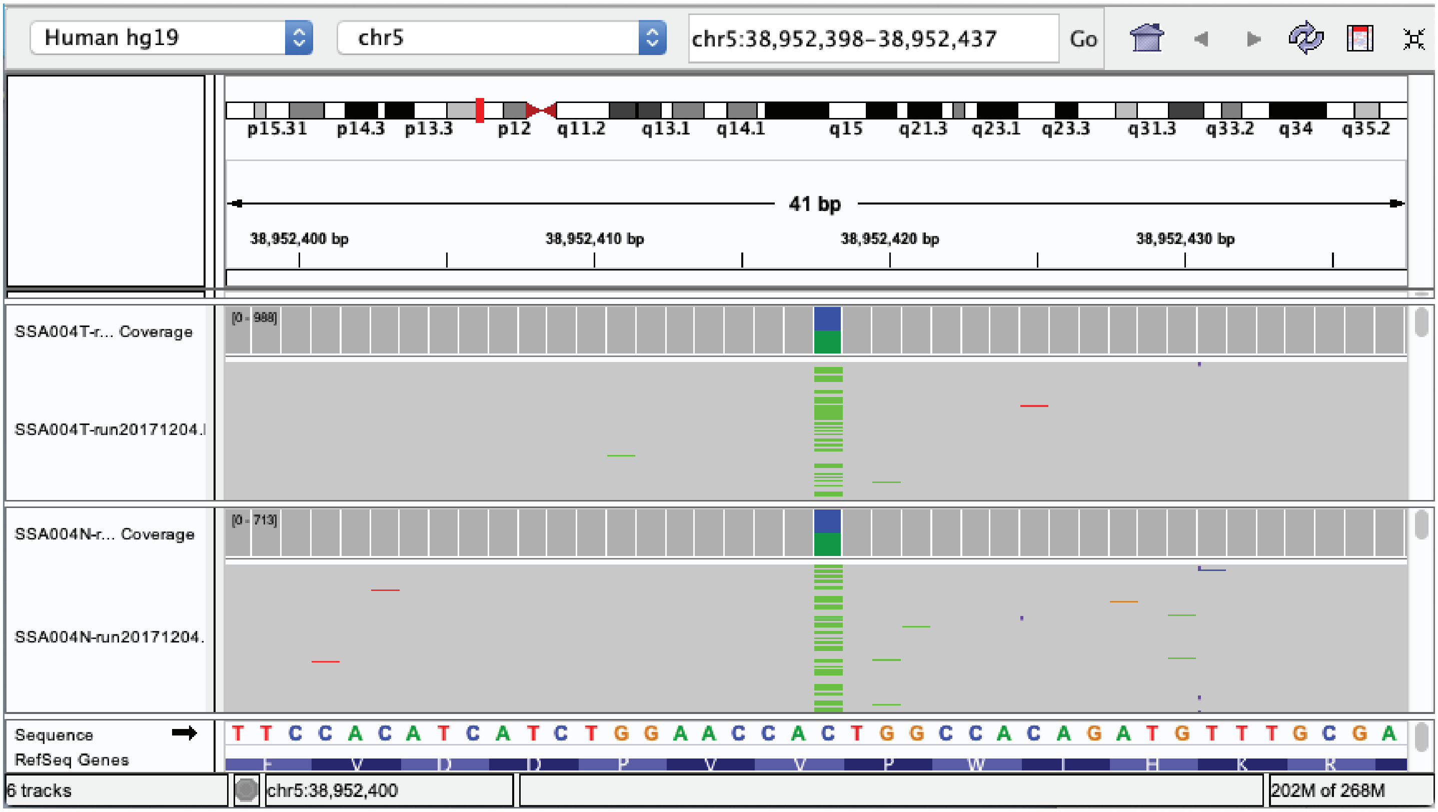


B

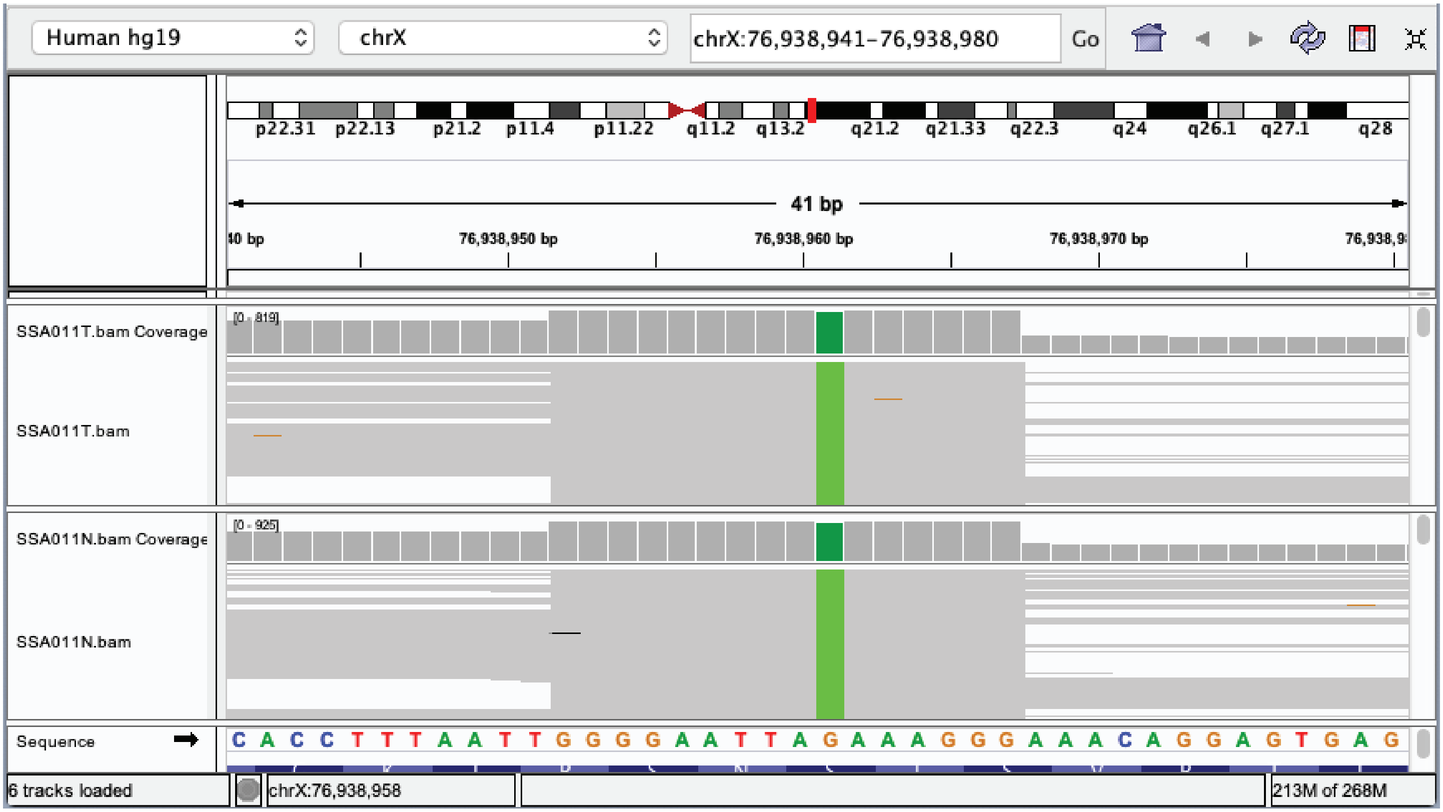

### Supplemental Figure 2

**Supplemental Figure 2**

**A**

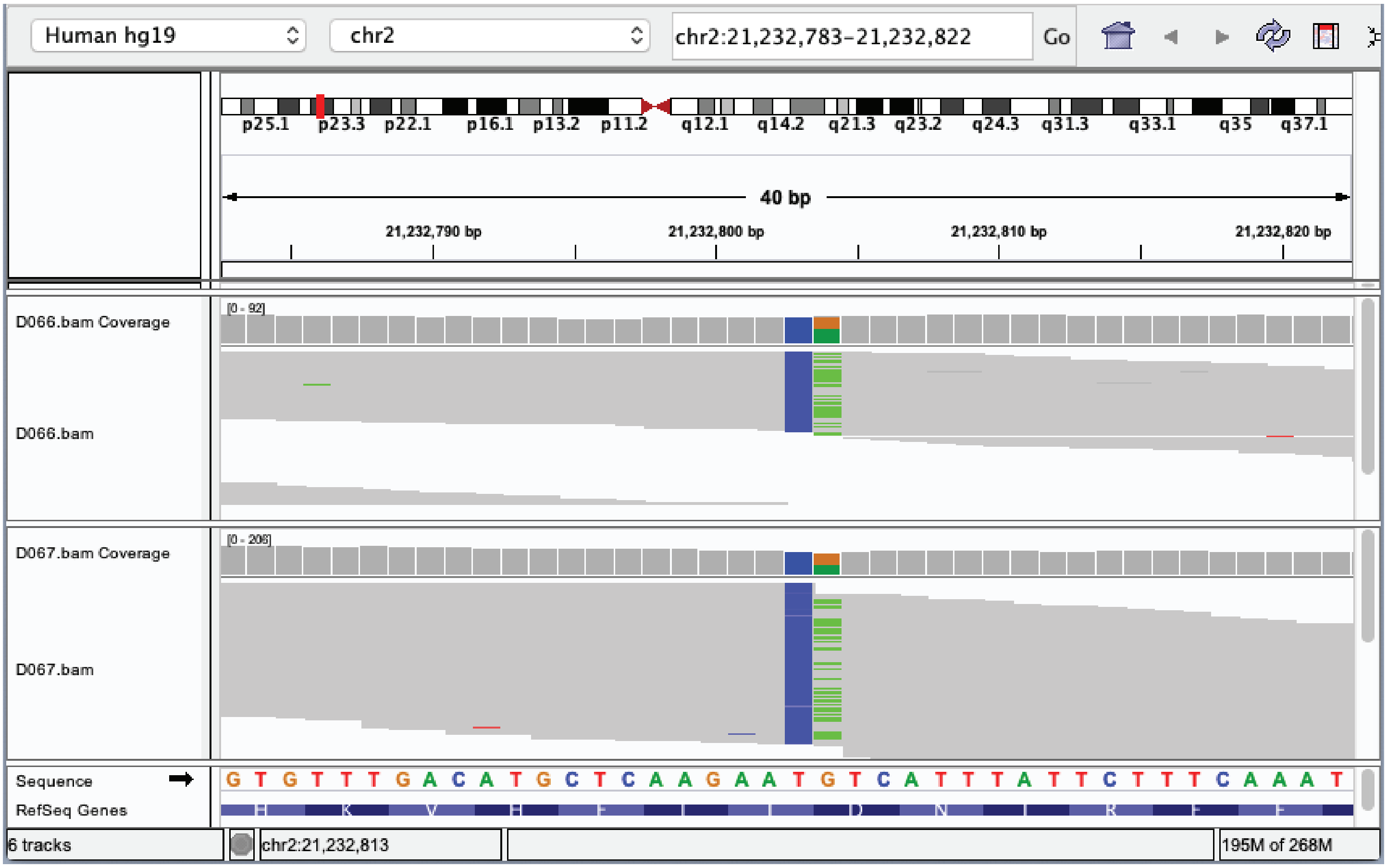


**B**

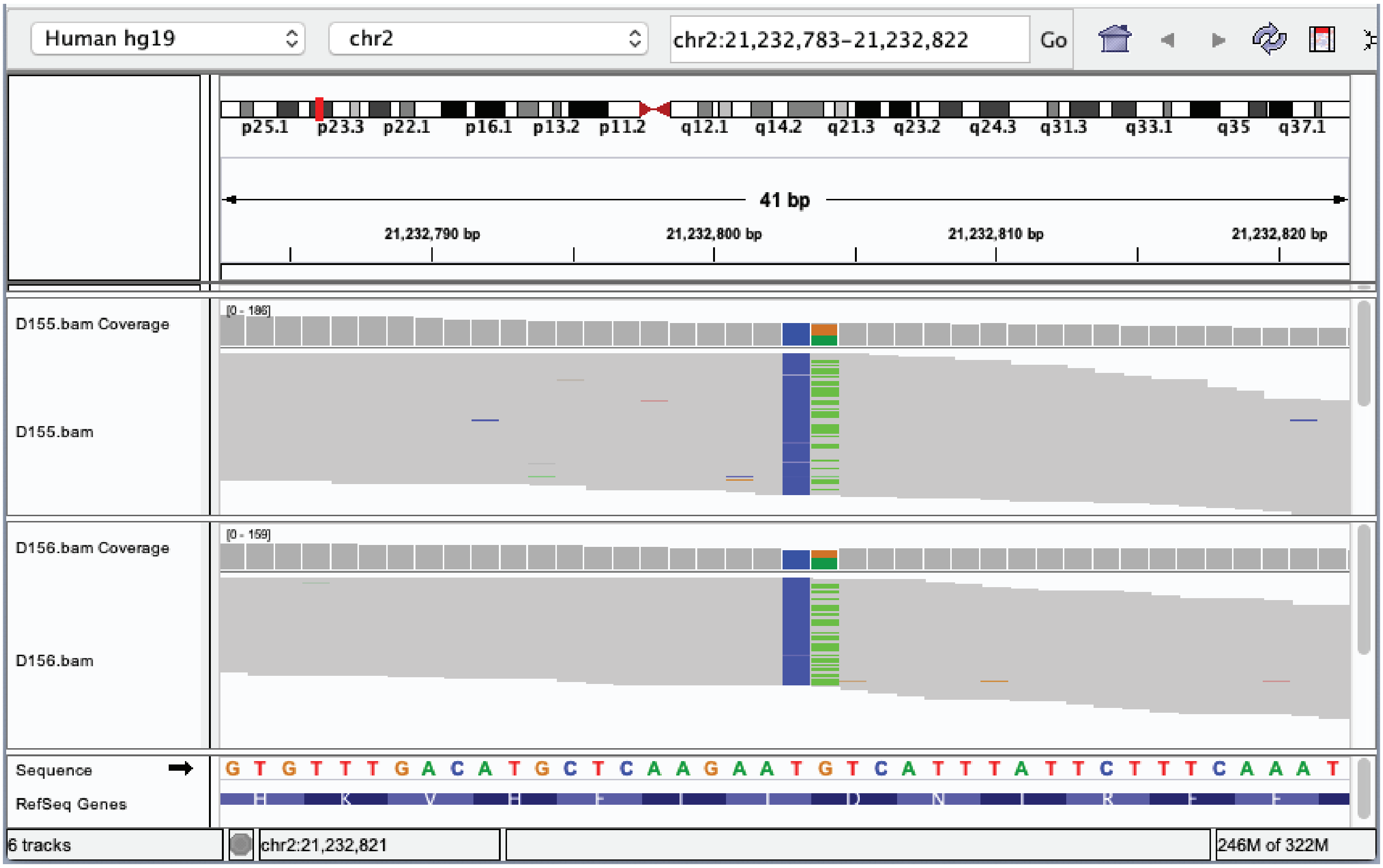
