## Supplementary legends for "PipeIT2: A tumor-only somatic variant calling workflow for Molecular Diagnostic Ion Torrent sequencing data"

**SUPPLEMENTAL FIGURE LEGENDS**

**Supplemental Figure 1:** Example of confirmed germline variants. Integrated Genome Viewer (IGV) view of two confirmed germline variants in the matched tumor and normal samples. The heterozygous SNP C>A at chromosome 5, position 38952418 in sample SSA004 and the homozygous SNP G>A at chromosome X, position 76938961 in sample SSA011.

**Supplemental Figure 2:** Example of confirmed germline variant *chr2:21232803:TG>CA* in orthogonal whole-exome sequencing data. IGV view of orthogonal whole-exome sequencing data, illustrating adjacent germline SNPs *rs584542* (*chr2:21232803:T>C*) and *rs1041968* (*chr2:21232804:G>A*) which were called as dinucleotide variant (DNV) *chr2:21232803:TG>CA* in *APOB*. Matched tumor (D067) and normal (D066) sample for patient HPU205 (upper panel) and matched tumor (D156) and normal (D155) sample for patient HPU212 (lower panel).
